# Murine aortic valve cell heterogeneity at birth

**DOI:** 10.1101/2024.07.22.604705

**Authors:** Theresa Bluemn, Julie R. Kessler, Andrew J Kim, Jenny Drnevich, Joy Lincoln

**Author notes:** Postnatal valve maturation. Corresponding Author: Joy Lincoln, 8701 Watertown Plank Road, Milwaukee, WI 53226.

## Abstract

Heart valve function requires a highly organized extracellular matrix (ECM) network that provides the necessary biomechanical properties needed to withstand pressure changes during each cardiac cycle. Lay down of the valve ECM begins during embryogenesis and continues throughout postnatal stages when it is remodeled into stratified layers and arranged according to blood flow. Alterations in this process can lead to dysfunction and if left untreated, heart failure. Despite this, the mechanisms that establish structure-function relationships of the valve, particularly during postnatal maturation, are poorly understood. To address this, single cell transcriptomics was performed on murine aortic valve structures at postnatal day one (PND1). Overall, 18 clusters of 7 diverse cell populations were identified, including a novel VEC subpopulation unique to PND1, and three previously unappreciated VIC subpopulations defined as “primitive”, “remodeling” and “bioactive”. Additional lineage tracing of the “primitive” VIC subpopulation in mice uncovered a temporal and spatial trajectory throughout postnatal maturation. In summary, this work highlights the heterogeneity of cell types within the aortic valve structure at birth that contribute to establishing and maintaining structure and function throughout life.

## INTRODUCTION

Heart valves are dynamic structures that adapt to ever-constant changes in the biomechanical environment to maintain unidirectional blood flow throughout life. This is largely achieved by a highly organized connective tissue consisting of layers of extracellular matrix (ECM) and diverse cell populations^1^. The valve ECM is organized into three layers arranged according to blood flow, and includes a collagen rich fibrosa layer for tensile strength, a ventricularis layer comprised of aligned elastin fibers for stretch, and a spongiosa layer of proteoglycans that absorb compressive stress^2^. In addition to the ECM, a layer of mechanosensitive valve endothelial cells (VECs) line the surface of the cusps, while valve interstitial cells (VICs) reside within and mediate deposition and turnover of the ECM in health and disease^3^. Previous studies have shown that molecular communication between VECs and VICs is needed to maintain ECM homeostasis, and therefore valve function^4–7^. In contrast, “VEC dysfunction” and “abnormal VIC activation” have been associated with “ECM disarray” and biomechanical weakness in valve disease^1,3^. Unfortunately to date, the mechanisms that underlie the pathological changes in the valve that lead to dysfunction are poorly understood, largely due to our lack of knowledge regarding how the valve ECM architecture is established and maintained in healthy individuals.

Formation of mature valve structures begins during embryogenesis and continues throughout postnatal life. For the aortic valve (AoV), the heart valve precursor cell pool is generated through a combination of endothelial-to-mesenchymal transformation (EMT) and neural crest cell migration into ECM-rich endocardial cushion structures. As development progresses, the cushions undergo remodeling and shaping to give rise to functional valve cusps and supporting structures by birth^8^. Postnatally, the cusps continue to elongate as the precursor cells lose their proliferative potential. By postnatal day 7 (PND7), single cell RNA-seq studies in mice report molecularly distinct VEC subpopulations, and ECM-producing VIC populations at this time^9^. In parallel, the ECM continues to undergo remodeling with enrichment of fibrillar collagens and proteoglycans^1,10^, and deposition of elastin fibers to on the ventricularis side^1^. As maturation progresses (PND30), the transcriptional profiles of VECs are largely unchanged from that at PND7, however the diversification of VIC subpopulations continue to evolve, which includes maturation into transcriptionally distinct antigen presenting, Tcf21, fibrosa, and matrifibrocytes subpopulations^9^. Studies in mice have shown that failure to form endocardial cushions with sufficient precursor cells during early, embryonic stages of valvulogenesis often leads to premature lethality before birth^11^. However, the regulators of postnatal maturation essential for completing functional valve formation are less well known.

Therefore, the goal of this current study was to use single cell transcriptomics to characterize AoV cell populations at PND1 in order to better understand valve maturation after birth and mechanisms of valve disease influenced by aberrations in this process. 14,289 cells were analyzed, from which seven broad cell populations, comprising 18 transcriptionally unique clusters, were identified based on gene expression. This includes a novel VEC subpopulation not previously described in the valve and three, dynamic VIC clusters that give rise to more mature VIC cell types later at later stages of maturation. This study provides the molecular profile of the PND1 AoV and describes the cellular heterogeneity needed for postnatal valve maturation and lifelong function.

## METHODS

A detailed description of the materials and methods used in this study can be found in the supplemental materials. The reported data have been deposited into GEO (accession ID: GSE269604).

## RESULTS

### Single nuclear gene expression profiling identifies cellular heterogeneity in the aortic valve structure at PND1

To define molecular and cellular heterogeneity of the healthy AoV structure at PND1, single nuclear RNA sequencing (10X Genomics) was performed in *C57/Bl6* mice^12^ (Fig. 1A). Three independent, pooled samples were sequenced, aggregated, and analyzed. Analysis by Seurat of a total of 14,289 cells identified 18 unique cell clusters (Fig. 1B) that were represented in all three replicate samples (Fig. S1A). The top 50 significant, upregulated cluster marker genes were assessed using Erichr, DAVID analysis and PanglaoDB to characterize the cell types representing each cluster^13–18^. Approximately 50% of the cells identified expressed cluster marker genes characteristic of the cell types traditionally found in the valve cusp based on published work^19^. Clusters 3, 7, 10, and 12 (2988 cells, 21%) exhibited enriched expression of *Cdh5* and were therefore classified as VECs, while clusters 4, 5, and 9 (3,317 cells, 23%) were enriched with *Col1a1* expression and identified as VICs. Cluster 11 (263 cells, 2%) exhibited strong expression of *Ptprc* indicative of immune cells, and cluster 18 (51 cells, 0.5%) was enriched for *Dct*, a marker of melanocytes. The remaining 50% of analyzed cells had transcriptional profiles consistent with cell types outside of the cusp and were therefore presumed to be within the aortic root structure. Clusters 1 and 15 (3579, 25%) were identified as smooth muscle cells based on known markers, including enrichment of *Acta2*. Clusters 2, 6, 8 and 14 (3661, 26%) were positive for *Tnnt2*, and therefore identified as cardiomyocytes. Clusters 13 and 16 (369, 3%) were enriched for *Cdh5* expression, however clustered independent from clusters 3, 7, 10 and 12. Finally, cluster 17 (61, 0.5%) cells expressed *Adipor2*, a marker of adipose cells (Fig. 1C, Fig. S1B). Single nuclear transcriptomics unveiled heterogeneous subpopulations within the seven major cell types identified based on their transcriptional diversity. The heterogeneity of the subpopulations comprising VECs (cluster 3, 7, 10, 12), VICs (cluster 4, 5, 9) and immune cells (cluster 11) were studied further based on their role in valve development, maintenance, and disease^20–22^.

**Figure 1.**
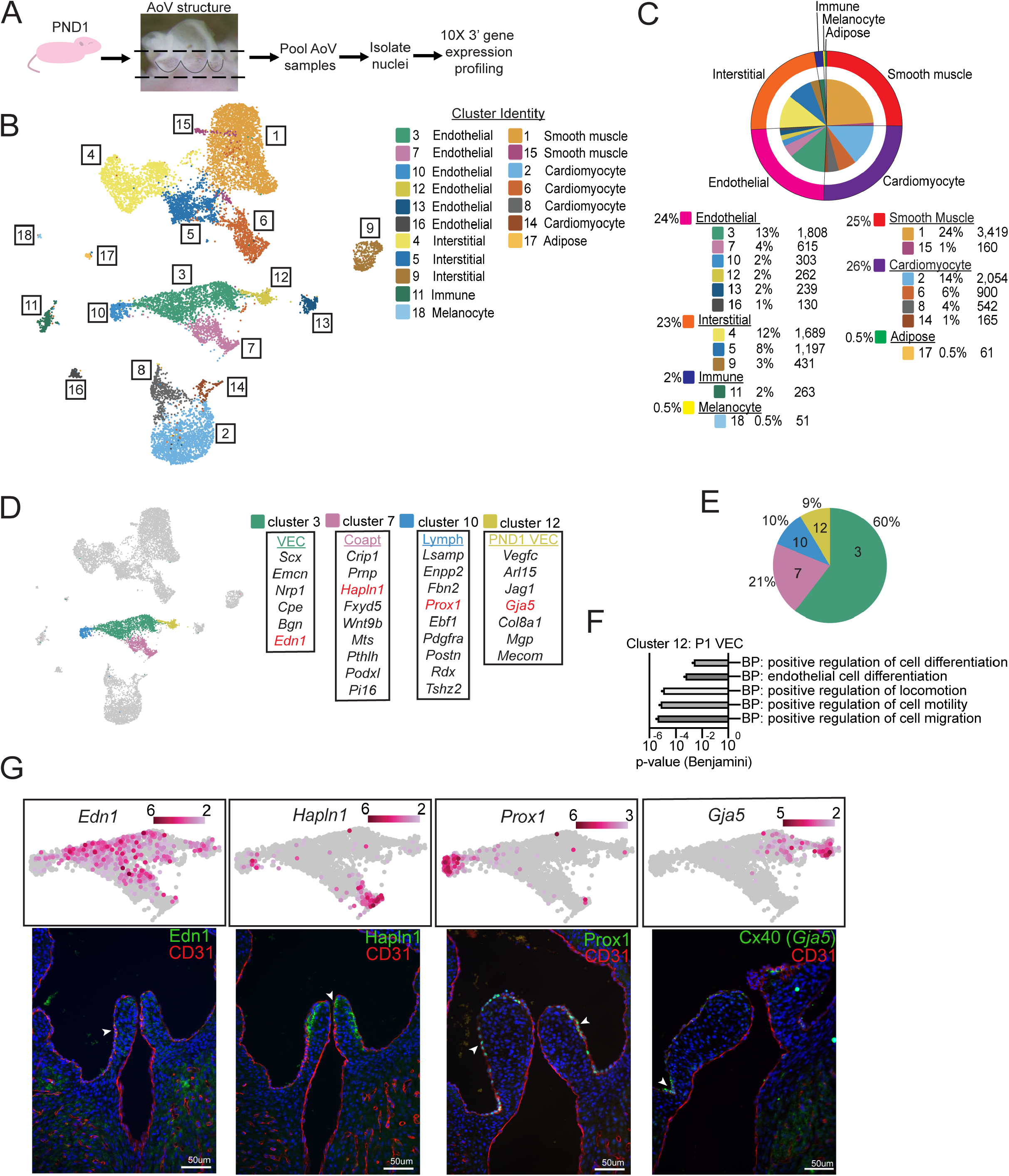
Single nuclear gene expression profile of the postnatal day 1 aortic valve structure. (A) Experimental scheme. Aortic valve (AoV) structures from *C57/Bl6* PND1 mice were processed for nuclei isolation and single nuclear gene expression profiling. (B) UMAP projection of 14,289 cells identifying 18 unique unsupervised graph-based gene expression Seurat clusters. Aggregated data of n=3 pooled aortic structures replicates, 22-24 aortic roots per replicate. (C) Pie chart depicting percentage of cells and cell number per cluster (inner circle) and per cell type (outer circle). (D) UMAP projection highlighting valve endothelial cells (cluster 3, 7, 10 and 12) based on published gene expression signatures. (E) Pie chart representing the percentage of cells identified in each VEC subpopulation. (F) DAVID analysis of GO-terms (Biological Processes) of the significantly, upregulated genes identified in cluster 12 compared to clusters 3, 7, and 10 (log_2_foldchange≤0.58; adj pvalue≤0.05). (G) (Top) Feature plots of UMAP projection of VEC subpopulation gene markers. Scale bar depicts log_2_fold-change. (Bottom) Representative immunofluorescence images of PND1 AoV structures identifying VEC subpopulations. Edn1: cluster 3 traditional VEC, Hapln1: cluster 7 coapt VEC, Prox1: cluster 10 lymph VEC, Gja5: cluster 12 PND1 VEC. CD31 to highlight the general endothelial cell population and DAPI (blue) staining to depict nuclei. Arrowhead highlights regions of localization of positive cells of cluster marker gene expression.

### Diversification of valve endothelial cell subpopulations at PND1

VECs serve as mechanosensitive signaling centers and additionally form an endothelial barrier over the cusp to prevent infiltration of circulating risk factors. Previous bulk RNA-seq studies from our lab indicated that VECs exhibit functional diversity from embryonic development through aging, despite few molecular changes^4^. Here, VEC clusters 3, 7, 10, and 12 were further investigated to characterize subpopulation identity at the single cell level by cross-referencing with published data performed at PND7 and PND30^9^. Interestingly, clusters 3, 7, and 10 shared similar gene expressions profiles to published VECs, including include *Edn1* (traditional VECs), *Hapln1* (coapt-VECs), *Prox1* (lymph-VECs), respectively (Fig. 1D). However, cluster 12 (9% of VEC population) showed a unique gene expression profile that includes *Gja5, Jag1* and *Mecom*, referred to as “PND1 VECs” (Fig. 1D, E). DAVID analysis of significantly upregulated genes in “PND1 VECs” compared to other VEC clusters 3, 7, and 10, unveiled association with “positive regulation of cell migration”, “positive regulation of cell motility”, “positive regulation of cell locomotion”, “positive regulation of cell differentiation” and “endothelial cell differentiation” (Fig. 1F). Immunohistological examination of the VEC clusters revealed, as expected,^9^ Hapln1^+^ coapt-VECs localized within the coaptation region of the cusp, Prox1^+^ lymph-VECs on the fibrosa side of the valve, and traditional VECs, (Edn1^+^) distributed over the AoV surface (Fig. 1G). Notably, Cx40^+^ (*Gja5*) expression, as an indicator of “PND1 VECs” was detected within the endothelium near the attachment zone hinge of the cusp (Fig. 1G). Overall, the majority of VEC subpopulations transcriptionally and spatially resemble that previously reported at PND7 and PND30, however a unique PND1 VEC subpopulation is regionally present.

### Tissue resident macrophages are the primary immune cell in the aortic valve at PND1

Immune cells play critical roles in pathological ECM remodeling, including myxomatous degeneration^23,24^. Here, cluster 11 was identified as a PND1-immune cell population based on enrichment of known macrophage gene markers (*Cx3cr1, Csf1r* and *Dab2*, Fig. S2A), and exhibited gene expression signatures associated with biological processes related to “myeloid cell homeostasis”, “myeloid cell development”, “positive regulation of myeloid cell differentiation”, “negative regulation of myeloid cell differentiation” and “macrophage differentiation” (Fig. S2B). Compared to all other cell clusters, *Mrc1*, a marker of tissue-resident macrophages^21^ was the most significantly enriched mRNA in cluster 11. To validate, immunofluorescence was used to detect CD45 (*Ptprc*) and CD206 (*Mrc1*) expression in sister sections of PND1 AoV structures and identified CD45^+^ cells throughout the valve cusp with regionalized overlap of CD206^+^ cells, suggesting tissue resident macrophages (Fig. S2C). Notably, Ccr2, a marker of bone marrow-derived circulating macrophages, did not meet the statistical thresholds of expression level and immunofluorescence was unable to detect protein expression (Fig. S2C). Gene signatures of innate and adaptive immune cells including, dendritic cells, mast cells, T-cells, B-cells or NK-cells, were not identified in cluster 11 compared to all other cells of the remaining 17 clusters. Taken together, this demonstrates that the primary immune cell type present in PND1 aortic valves is CD45^+^ CD206^+^ tissue-resident macrophages.

### The aortic valve interstitial cell population is highly heterogenous at PND1

VIC clusters 4, 5 and 9 were further examined to assess subpopulation identity. Figure 2A shows that VIC clusters 4, 5, and 9 exhibit comparatively unique gene expression profiles at PND1. VICs within cluster 4 associate with transcription factors commonly associated with post-EMT valve precursor cells within the endocardial cushion including *Msx1, Sox9, Tbx20* and *Scx*^25–28^ (Fig. 2A) and comprise approximately 50% of all PND1 VICs (Fig. 2B). DAVID analysis of the significantly upregulated genes in cluster 4 compared to clusters 5 and 9, show gene ontology (GO) terms of “positive regulation of cell migration”, “positive regulation of motility”, “cell differentiation”, “mesenchymal cell differentiation”, and “cell proliferation” (Fig. 2C). Based on this, cluster 4 are considered “primitive” VICs. Spatially, primitive-VIC localize to the hinge region (Fig. 2D), and consistent with GO, reside in the same region as actively cycling cells (Fig. S3A) and appear to migrate throughout the cusp by PND14 (Fig. S3B).

**Figure 2.**
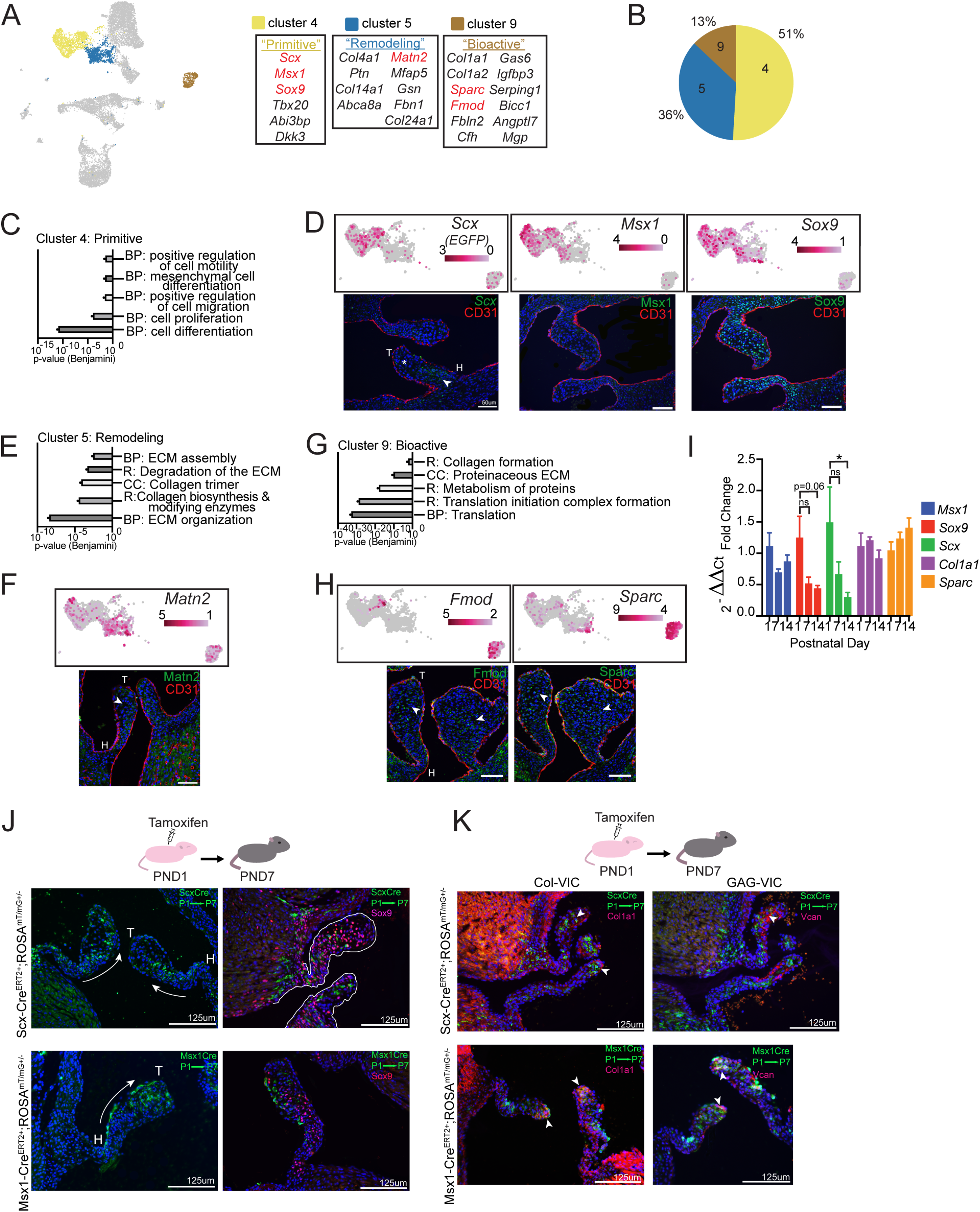
Valve interstitial cell profiling in the murine aortic valve at postnatal day 1. (A) UMAP projection highlighting VICs (cluster 4, 5, and 9) with gene expression signatures of each subpopulation. (B) Pie chart representing the percentage of cells identified in each VIC subpopulation. (C) DAVID analysis of GO-terms of the significantly upregulated genes identified in cluster 4 compared to clusters 5 and 9 (log_2_foldchange≤0.58; adj pvalue≤0.05). (D) (Top) Feature plots of UMAP projection of primitive-VIC subpopulation (cluster 4) gene markers *Scx (EGFP), Msx1* and *Sox9*. Scale bar depicts log_2_fold-change. (Bottom) Representative immunofluorescence images of primitive-VIC gene markers, Sox9 and Msx1 protein and *Scx* RNA transcript in AoV structures throughout valve maturation at PDN1. (E) DAVID analysis of GO-terms of the significantly, upregulated genes identified in cluster 5 VIC subpopulation cluster compared to clusters 4 and 9 VIC subpopulations (log_2_foldchange≤0.58; adj pvalue≤0.05). (F) (Top) Feature plot of UMAP projection of remodeling-VIC subpopulation (cluster 5) gene marker *Matn2*. (Bottom) Representative immunofluorescence images of Matn2 staining in the PND1 AoV structure. (G) DAVID analysis of GO-terms of the significantly, upregulated genes identified in cluster 9 VIC subpopulation cluster compared to cluster 4 and 5 VIC subpopulations (log_2_foldchange≤0.58; adj pvalue≤0.05). (H) (top) Feature plot of UMAP projection of bioactive-VIC subpopulation (cluster 9) gene markers *Fmod* and *Sparc*. (bottom) Representative immunofluorescence images of Fmod and Sparc staining in the PND1 AoV structure. (I) RT-qPCR of VIC subpopulation gene markers during postnatal valve maturation. n=3-7 individual samples of n=1-6 pooled aortic structures. *pvalve≤0.05 unpaired, nonparametric Mann-Whittney t-test. (J) Lineage tracing of cluster 4 primitive VIC subpopulation during valve maturation in *Scx-Cre*^*ERT2*^ and *Msx1-Cre*^*ERT2*^ mice bred with Rosa^mT/mG^. *Scx-Cre*^*ERT2*^*;Rosa*^*mT/mG*^ (Top) and *Msx1-Cre*^*ERT2*^*;Rosa*^*mT/mG*^ (Bottom) mice were injected with tamoxifen at PND1 and harvested at PND7. (Left) Immunofluorescent images to detect GFP as an indication of *Cre-*recombination. Arrows indicate trajectory of GFP^+^ cells. (Right) Immunofluorescent images of GFP immunoreactivity to depict cluster 4 primitive VIC subpopulation cell trajectory over time with co-staining of Sox9 (primitive-VIC marker). (K) Immunofluorescence images of cluster 4 primitive VIC subpopulation lineage tracing from PND1 to PND7 (as performed in J) co-stained with Col-VIC (Col1a1) and GAG-VIC (Vcan) population markers. CD31 identifies endothelial cells, and DAPI staining depicts nuclei. Scale bar depicts 50μm. BP: biological process, CC: cellular component, R: Reactome, H: valve hinge region, T: valve tip region, arrowhead highlights regions of localization of positive cells of cluster marker gene expression, star highlights region of localization of negative cells of cluster marker gene expression.

The gene expression profile of cluster 5 shows upregulation of ECM genes including *Col4a1, Col14a1* and *Matn* ^29^ (Fig. 2A) and comprise 36% of all VICs at PND1 (Fig. 2B). GO and Reactome pathway analysis of the significantly upregulated genes in cluster 5 compared to clusters 4 and 9, associate with “ECM assembly”, “ECM organization”, “collagen biosynthesis and modifying enzymes”, “collagen trimer” and “degradation of the ECM” (Fig. 2E). As a result, this VIC subpopulation is referred to as “remodeling” VICs, and by immunofluorescence for Matn2, exhibit dispersion throughout the cusp at PND1 (Fig. 2F). Finally, cluster 9 VICs are enriched for *Sparc* and *Fmod*, in addition to *Col1a1* (Fig. 2A) and appear to be the least abundant VIC subpopulation at PND1 (13%) (Fig. 2B). GO profiling of cluster 9, compared to clusters 4 and 5, suggests roles in “collagen formation”, “proteinaceous ECM” and biosynthetic processes including “metabolism of proteins”, “translation” and “translation initiation complex formation” (Fig. 2G) and therefore were considered “bioactive” VICs. Sparc^+^,Fmod^+^ bioactive-VICs reside in similar regions of the cusp tip and belly region as remodeling-VICs (Fig. 2H). By qPCR analysis, it appears that during postnatal maturation, primitive-VIC gene markers (*Sox9, Scx, Msx2*) decrease over time, while markers of bioactive-VICs (*Col1a*1, *Sparc*) increase at PND7 and PND14 (Fig. 2I). Worthy of mention, the three PND1 VIC subpopulations did not share significant molecular overlap with previously reported transcriptional profiles at PND7 nor PND30^9^, suggesting that these novel subpopulations that warrant further investigation.

### Primitive valve interstitial cells are present throughout postnatal maturation

As the cluster 4 primitive-VICs comprise the largest VIC subpopulation at PND1, lineage tracing studies were performed throughout postnatal maturation to determine their fate. For this, *Msx1-Cre*^*ERT2*^ and *Scx-Cre*^*ERT2*^ bred with *ROSA*^*mT/mG*^ reporter mice were treated with one dose of tamoxifen to activate *Cre-*recombinase at PND1^30–32^, and analysis was performed at PND7 (Fig. 2J) and PND14 (Fig. S3C). As shown in Fig. 2D, both *Msx1* and *Scx* lineages were localized within the hinge region at PND1, however, by PND7, GFP^+^ cells indicating Cre recombination, progressed towards the tip region of the cusp (Fig. 2J) resulting in widespread distribution at PND14 (Fig. S3C). Interestingly over this time, a subset of the GFP^+^ *Cre-*lineages maintain their primitive-VIC molecular identity as indicated by co-expression of GFP^+^ with Sox9^+^ at PND7 and PND14 (Fig. 2J, Fig. S3C), while other GFP^+^ cells co-express markers of previously described ECM-VIC subpopulations at PND7^9^, including Col1a1^+^ (Col-VICs) and Vcan^+^ (GAG-VICs)^9^ within the tip region (Fig. 2K). Therefore, suggesting that a sub-population of primitive-VIC lineage remain “immature”, while others take on more mature VIC markers at later stages of postnatal maturation, and this is spatially dependent.

## DISCUSSION

The formation of fully functioning heart valve structures requires the tight regulation of molecular and cellular processes beginning during embryonic development and continuing throughout postnatal maturation until the stratified ECM layers become distinguished and valve cells reduce proliferation and undergo further diversification^1,9,20^. While embryonic valvulogenesis has been well-studied, the mechanisms underlying maturation after birth are poorly understood, yet are likely critical for completing valve formation in preparation for lifelong function. Therefore, the goal was to determine the heterogeneity of valve cells present at birth that serve as the “pool” for postnatal maturation of the AoV structures through transcriptomic single cell sequencing. We show that murine AoV structures are comprised of several molecularly diverse cell clusters including VIC and VEC subpopulations that contribute to valve maturation after birth.

Three of the four VECs subpopulations identified here were molecularly comparable to that published at PND7 and PND30, which is surprising given the drastic change in the hemodynamic environment that mechanosensitive VECs experience during postnatal maturation as the patent foramen ovale begins to close. Although the molecular homology of VECs during the postnatal period was similarly observed in previous bulk RNA-seq studies spanning embryonic, postnatal and adult life^4^. Together, suggesting that VECs remain molecularly homeostatic throughout life in the absence of disease. Despite similarities, this study did identify a unique VEC subpopulation termed “PND1-VECs” that were enriched for *Gja5, Jag1* and *Mecom* expression and comprised ∼9% of total VECs at PND1. These cells are localized within the hinge region of the AoV cusp close to the point of attachment and based on their gene expression profile associated with migration, motility and differentiation. It is speculated that this subpopulation has a role in growth and elongation of the cups, and could potentially be remnants of embryonic development, and/or required for early postnatal maturation (as they are not observed by PND7). Global knockouts of either, *Jag1* or *Mecom* result in embryonic lethality by day E10.5 due to defects in cardiac morphogenesis^33–35^, and therefore valve-specific studies are required to gain further insights into their roles in postnatal valve maturation.

In contrast to the VEC population, there is dynamic diversification within the PND1 VIC population. We identified at least three PND1 VIC subpopulations and based on mRNA expression and associated GO analysis, refer to them as “primitive-VICs” (*Sox9, Msx1, Tbx20, etc*.), “remodeling-VICs” (*Ptn, Matn2, Col4a1, etc*.), and “bioactive-VICs” (*Fmod, Sparc, Mgp, Col1a1, Col1a2, etc*.). Each PND1 VIC subpopulation is transcriptionally unique from subpopulations identified at later postnatal timepoints^9^, suggesting that VICs are temporally dynamic and adaptive. Remodeling-VICs comprise one third of the total VIC population present at birth while bioactive-VIC comprise the smallest percentage (13%), and while both these VIC subpopulations express a variety of ECM-related mRNAs including proteoglycans, collagens and remodeling enzymes, they are molecularly distinguishable. For example, remodeling-VICs are more highly enriched with collagen, while bioactive-VICs additionally exhibit enrichment in metabolic processes and translation gene signatures. The role of these VICs is not clear, but the upregulation of translation and metabolic-associated mRNAs by bioactive-VIC could be a hallmark of high activity that includes differentiation and proliferation^36,37^, in contrast to remodeling-VICs that are more quiescent, but highly secretory. Notably, we did not identify a singular ECM producing VIC at PND1, unlike at PND7 that includes Col-VICs and GAG-VICs^9^, which suggests that compartmentalized ECM production likely occurs after PND1.

Primitive-VICs are the largest contributor to the total VIC population at PND1 (51%) and express mesenchyme cell transcription factors. At birth, these cells localize within the region close to the hinge of the cusps, whereas remodeling-VICs and bioactive-VICs are more widely distributed within the belly of the cusp. Then during postnatal maturation, the primitive-VIC expression pattern extends towards the tip within regions occupied by bioactive-VICs and remodeling-VICs by PND7. We therefore speculate that primitive-VICs are the least mature VIC subpopulation, and possibly remaining ‘valve precursor cells’ from the cushion, while remodeling-VICs and bioactive-VICs are more mature and participate in ECM remodeling and layer stratification during the first week of life. Together, this study further contributes to defining the hierarchy of VIC maturation, yet the plasticity and adaptability of this dynamic cell population in health and disease remains understudied.

Our study focused on characterizing physiologically normal subpopulations of AoV cells at birth and identified previously unappreciated clusters of VECs and VICs based on mRNA profiles. Now that potential regulators of postnatal maturation have been identified in a cell-specific manner, future studies can take targeted approaches to delineate the mechanisms that when altered, may give rise to valve pathology that manifests after birth.

## Acknowledgements

We thank the DNA Services Lab and High-Performance Biological Computing at the Roy J Carver Biotechnology Center, University of Illinois at Urbana-Champaign for assistance in the single nuclear gene expression study.

